# In-silico analysis of MHC genes in hereditary colorectal cancer shows identical by state SNP sharing affecting HLA-DQB1 binding groove

**DOI:** 10.1101/040436

**Authors:** Mahmoud E. Koko, Suleiman H. Suleiman, Mohammed O.E. Abdallah, Muhallab Saad, Muntaser E. Ibrahim

## Abstract

**Background:** The role of Human Leukocyte Antigen (HLA) alleles in colorectal cancer susceptibility, development and progression is the focus of ongoing scrutiny. MHC polymorphisms in a Sudanese family with hereditary colorectal cancer were studied using an *in silico* approach and the results were verified using The Cancer Genome Atlas (TCGA). In this family study, we tested for sharing of nucleotide polymorphisms identified by whole exome capture in *major histocompatibility complex* region and carried out *in-silico* prediction of their effects in tumor and control samples. SNPs were analyzed to highlight identical by state sharing, to identify runs of homozygosity, as well as to predict structural and functional effects using homology modeling, damaging effect predictions, and regulatory changes prediction.

**Results:** MHC II area showed significantly high degree of homozygosity in tumor samples. Non-synonymous SNPs shared *identical by state* (IBS) between tumor samples were predicted to affect HLA-DQB1 binding groove. A similar haplotype of these SNPs was identified in a TCGA colonic adenocarcinoma tumor sample. No significant regulatory effects (in the form of transcription factor or miRNA binding site variants) were predicted.

**Conclusions:** The results demonstrate IBS SNP sharing of markers affecting HLA-DQB1 binding specificity and probable loss of heterozygosity in MHC II region in colorectal cancer. The significance of this sharing in cancer pathogenesis remains to be established.

## INTRODUCTION

Colorectal tumors range from benign growths to invasive cancer and are predominantly epithelial-derived tumors (i.e., adenomas or adenocarcinomas). Colorectal carcinoma (CRC) is a leading cause of death in the world. Genome-wide analysis of gene mutations in CRC has identified acquired somatic mutations in several genes, highlighting the heterogeneity of the disease. In addition, each tumor usually shows a distinct mutational gene signature [1,2]. A portion of colorectal cancers display microsatellite instability [MSI; 15% of CRCs) characterized by defective DNA mismatch repair (MMR) system [3,4]. Genetic susceptibility to colorectal cancer is described primarily in familial colorectal cancer syndromes like hereditary nonpolyposis colorectal cancer (HNPCC). Notably, none of CRC susceptibility genetic polymorphisms were found in the *major histocompatibility complex* (MHC) region located in chromosome 6 (6p21.3), which is one of the most polymorphic areas in the human genome. It has been argued that *human leukocyte antigens* (HLA) polymorphisms can explain much of the diversity in cancer predisposition, progression and prognosis.

The loss of HLA gene expression owing to viral infection, somatic mutations or other causes may have important effects on immune suppression and cancer development. It is well known that some infection-related malignancies (e.g. Epstein-Barr virus, Human Papilloma Virus) show HLA-related predisposition or protection. Bernel–Silva et al. found that HLA-DRB1*14 allele seems to confer protection against cervical cancer [5]. However, other types of cancer with no clear relation to infectious agents show HLA predisposition as well. Although large genome-wide association studies have failed to demonstrate association between colorectal cancer and HLA SNPs, a previous study reported that HLA-DQA1* 0201 was significantly less common in patients with colonic carcinoma than controls [6]. HLA DQB*03032 and HLA DRB1*11 alleles may have a protective role in human breast cancer. HLA DRB1*11 alleles were found significantly overrepresented (P<0.0001) in controls as compared with patients with early-onset breast cancer [7]. HLA DRB1*0901 allele was found to be more prevalent in the patients of esophageal carcinoma compared to healthy controls in Hubei Han Chinese, possibly indicating higher susceptibility [8]. El-chennawi et al. found a significantly increased frequency of HLA-DRB1*04, DRB1*07, and DQB1*02 in Hepato-Cellular Carcinoma Egyptian patients versus control group, as well as a significantly decreased frequency of DQB1*06 and DRB1*15 [9]. It was shown that somatic mutations in cancer can affect the binding groove of HLA genes, possibly altering the binding specificities of these molecules. Variants showing potential epigenetic, peptide-loading function and T-cell immune response were correlated with the effects of HLA-A*11:01, a protective HLA-A allele against nasopharingeal carcinoma in Malaysian Chinese population. Most other HLA-A variants did not appear to possess any potential function [10].

This study aimed to characterize MHC SNPs in HNPCC by analyzing MHC polymorphisms in a Sudanese family with hereditary colorectal cancer. Analysis of exomes of tumor and control samples from this family showed an interesting shared mutation pattern in tumor samples as described by Suleiman et al. [11]. In this study, we focused on MHC genes and found features of identical by state SNP sharing affecting HLA-DQ binding groove and thus the binding specificity.

## METHODS

### Study Materials and Exome Sequencing

The study examined exome sequences from an extended Sudanese family with a history of hereditary colorectal cancer previously described by Suleiman et al. [11]. We took two cancer patients and two related controls. DNA was extracted using standard techniques from tumor samples obtained following surgical resection from the two cancer patients and control blood samples from two controls (a sister to one patient and another distant cousin from a branch of the extended family). Quality of extracted DNA was assured and whole exome sequencing was performed (BGI©, China) on Illumina HiSeq2000 platform (Illumina, USA). Alignment was performed using BWA [12]. SNP discovery and genotype calling was done using SOAPsnp [13]. INDEL calling was performed using ATLAS2 [14]. This study utilized the results of called SNPs in MHC area on chromosome 6 by filtering MHC genes from the total variant calls.

### Variants Analysis

SNPs found in the MHC region in study samples were studied in terms of reported association to cancer, amount of variant sharing, homozygosity and variants effect prediction. SNPs number, sharing, identity, heterozygozity, predicted structural/functional outcomes of observed variants, possible effect on regulatory sites, as well as haplotype sharing was studied using In-Silico tools and databases as detailed below.

### Annotation and Effect Prediction

The identified SNPs were annotated using SNPNexus tool [15]. Variants were tested for possible Transcription Factor Binding Sites (TFBS), miRNA binding using SNPNexus. Genomic Evolutionary Rate Profiling (GERP++) conservation scores [16] were used to assess if mutations affected conserved sequences of genome. Relation of SNPs to Sequence Feature Variant Types sites (SFVTs) were checked using definitions on NCBI protein database. NCBI Genetic Association Database [17] and Phenotype-Genotype Integrator [18] were searched for reported SNPs in our samples. Damaging effects of non-synonymous coding variants were predicted using Consensus Deleteriousness (ConDel) scores [19]. iMutant2 [20] was used to predict changes in protein stability caused by single mutations with high probability of damaging effect reflected in predicted free energy change (DDG) scores. Swiss-Model [21] was used to predict the 3D structure of mutated HLA proteins using reference UniProtKB sequences. UCSF Chimera [22] was used to visualize modeled PDB files. Regulatory effects of SNPs were assessed using multiple tools: RegulomeDB scores [23], Alibaba 2.1 [24] and TFSEARCH [24].

### Marker sharing

Beagle/fastIBD [25] software package was used for Identity By Descent (IBD) and Homozygosity By Descent (HBD) inference. Significance of marker sharing was tested by Chi-square test at 1000 Monte Carlo permutations using Haploview [26]. Mantel statistic was used to test haplotype sharing among individuals using Step-down adjusted p-values (adjusting for the Family Wise Error Rate) calculated at 1000 Monte Carlo Permutations using the software Tomcat v1.0 [27].

### Homozygosity

proportions of homozygous SNPs out of total markers seen in each sample were calculated. Chi square test with Monte Carlo simulations and Marascuilo procedure (using Microsoft^®^ Excel 2010/XLSTAT v.2014.2.07, Addinsoft Inc., USA) was used to test significance of difference between multiple proportions. Runs of Homozygosity (ROH) were estimated depending on the number of consecutive homozygous markers. Siraj et al. [28] defined a Run of Homozygosity in colorectal cancer patients as genomic regions where minimum of fifty consecutive SNPs were homozygous using SNP arrays. However, because only roughly about 30% of SNPs captured in human genome are found in areas targeted by exome sequencing [29], a run of 17 consecutive homozygous SNPs in exome sequences would be expected to indicate homozygous genomic region. A rather arbitrary threshold of 20 SNPs was set here as an indicator for the presence of Run of Homozygosity.

## RESULTS

A total of 77 single nucleotide variants were seen in four samples (tumor samples P17 & P61; control samples P26 & P39). Twenty seven of them were exonic SNPs while fifty two were found in intronic regions. Those variants were located on one MHC I gene (HLA-A), and four MHC II genes (HLA-DRB1, HLA-DRB5, HLA-DQA1 and HLA-DQB1). No variants were found in MHC III area. Only one intronic variant located in HLA-DQB1 gene was novel (g.32628745G>T). It was seen in control sample P26. Table (1) summarizes SNPs found in HLA genes. Apart from an exonic deletion of HLA-G gene (rs41557518) affecting one tumor sample (P17) there were no other exonic deletions in MHC area. Majority of exonic SNPs lay in evolutionary conserved areas of MHC II containing HLA-DQA1 & HLA-DQB1 genes sequences. Figure (1) show MHC II region SNPs positions and sharing between study samples. Figure (2) outlines SNVs sharing at four MHC II genes harboring variations. An area between g.32629739 and g.32630014 with high conservation score (GERP++ RS score 596) contained SNPs unique to control samples as well as shared SNPs among samples. The shared SNPs between CRC tumors samples concentrated around conserved area in chromosome 6 between g.32632609 and g.32632854 (GERP++ RS score 288). Called SNPs did not affect sites identified as sequence features at NCBI protein database. HLA-A showed variations in tumor samples only. Details of SNVs positions seen in each sample are provided in supplementary (1).

**Table (1).** Tables. Single nucleotide variations detected in HLA genes in study samples. Variations are reposted as dbSNP rs IDs except for one novel variant (reported as hg19 genomic position and alleles).

| Gene | Sample | SNVs |
| --- | --- | --- |
| HLA-A | P17 | rs63225861 |
|  | P61 | rs1059449; rs41541222; rs63225861; rs14288; rs2975057 |
| HLA-DRB5 | P17 | rs3205665 |
|  | P39 | rs113965088; rs41546317 |
|  | P26 | rs1060153; rs3200923; rs3205665; rs113965088; rs41546317 |
| HLA-DRB1 | P17 | rs1064701; rs9269693; rs35985423 |
|  | P61 | rs9269693; rs35985423 |
|  | P39 | rs9269693 |
|  | P26 | rs142078339; rs1064701; rs12363; rs1064697; rs9269693; rs35985423; rs143082263 |
| HLA-DQA1 | P17 | rs28437906; rs9272544; rs10093; rs4467816; rs9272851; rs8227; rs112931520; rs7143; rs28376734; rs28538060 |
|  | P61 | rs28636516; rs28437906; rs9272544; rs10093; rs4467816; rs9272851; rs8227; rs112931520; rs7143; rs28376734; rs28538060 |
|  | P39 | rs28435909; rs28636516; rs28437906; rs9272544; rs12722051; rs10093; rs36219699; rs4467816; rs9272718; rs35975014; rs62404092; rs1130116; rs36027005; rs9272851; rs7708; rs8227; rs112931520; rs77015062; rs7143; rs9492; rs28376734; rs36216730; rs28538060 |
|  | P26 | rs28435909; rs28437906; rs9272544; rs12722051; rs10093; rs36219699; rs4467816; rs9272718; rs35975014; rs62404092; rs1130116; rs36027005; rs9272851; rs7708; rs8227; rs112931520; rs77015062; rs7143; rs9492; rs28376734; rs36216730; rs28538060 |
| HLA-DQB1 | P17 | rs17412886; rs9273671; rs9273677; rs9273713; rs9273723; rs36233043; rs9273912; rs1049107; rs1130397; rs17412833; rs1130392; rs1130390; rs1071637; rs1063318; rs1049069; rs9274526 |
|  | P61 | rs17412886; rs9273671; rs9273677; rs9273713; rs9273723; rs9273912; rs1049107; rs17412833; rs1130392; rs1130390; rs1071637; rs1063318; rs1049059; rs9274526 |
|  | P39 | rs1071639; rs17416274; rs28688207; rs17412886; rs9273671; rs9273713; rs117780937; rs36233043; rs9273912; rs75643968; rs701564; rs1130397; rs17412833; rs1130392; rs1049056; rs1049055; rs1049053; rs9274526 |
|  | P26 | rs1071639; rs28688207; rs117988605; rs117867884; g.32628745G>T; rs9273534; rs2854273; rs9273677; rs9273713; rs117780937; rs36233043; rs9273912; rs75643968; rs701564; rs1049133; rs1049130; rs1049088; rs1049107; rs28746795; rs1130397; rs17412833; rs1130392; rs1049059; rs67147537; rs9274526 |

**Figure (1).**
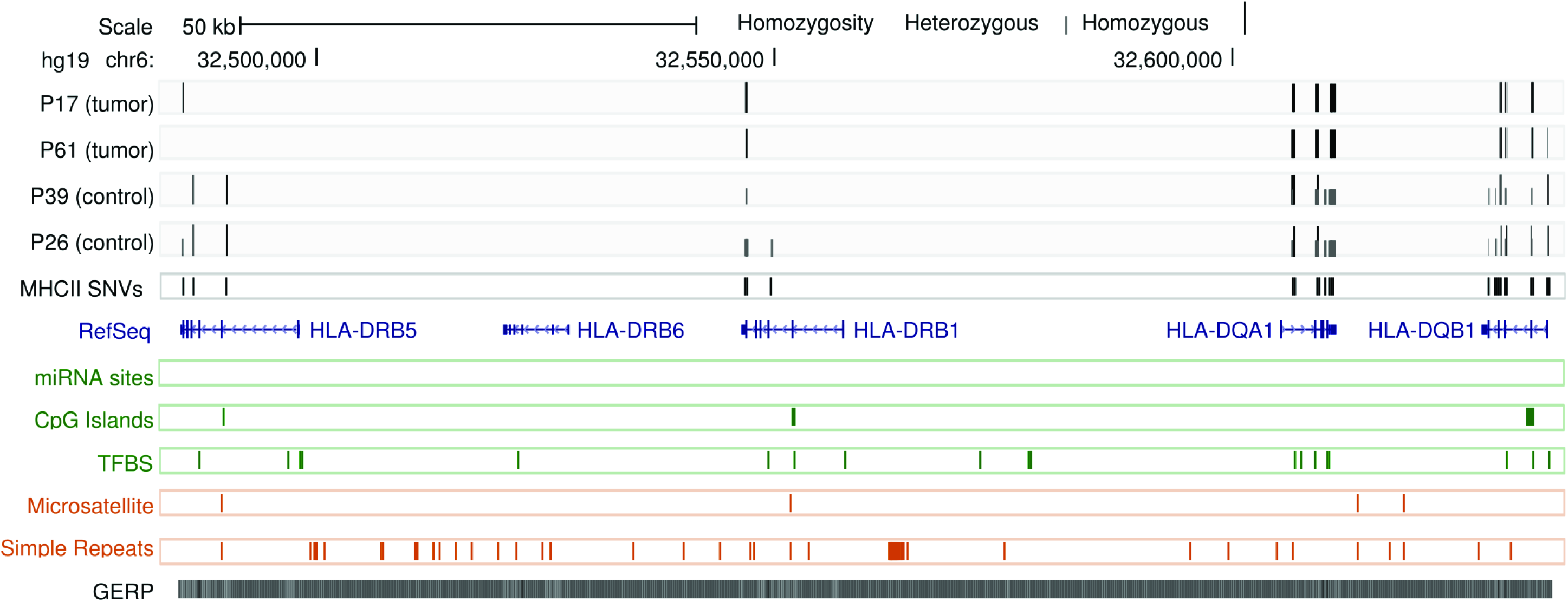
Position and homozygosity of SNVs in major histocompatibility II region in study samples. For each sample, variants are represented by short or long lines according to homozygosity. Annotation of MHCII area are show as well: RefSeq genes, TargetScan miRNA binding sites, conserved transcription factor binding sites (TFBS), CpG islands, microsatellites (with di/trinucleotide repeats), simple repeats, and genomic evolutionary rate profiling scores (as shading gradients). These annotations are obtained with modification from UCSC genome browser [43].

**Figure (2).**
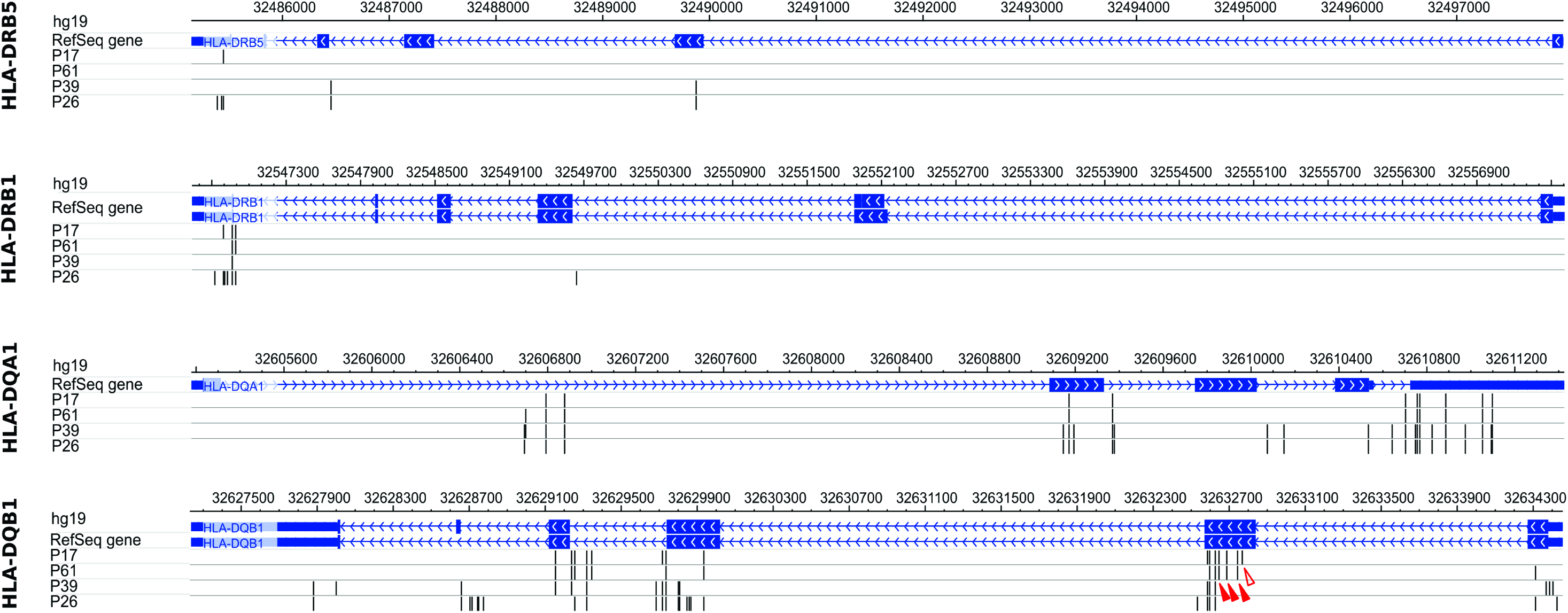
Sharing of SNVs in MHC II genes between study samples. Variants are represented as vertical lines according to hg19 genomic coordinates. Four nonsynonymous exonic variants affecting HLA-DQB1 are seen in tumors only (arrow heads); three of them are present in both cases (solid arrow heads) as well as in a TCGA sample (TCGA ID: TCGA-G4-6298) indicating that they are identical by state. This image was produced with modification using Wash U Epigenome Browser [44].

### Comparing polymorphisms between samples

The average number of SNPs per haplotype in each sample was 31, 31.5, 36.5, and 29 in samples P17, P61, P26 and P39, respectively. Sixteen SNPs were shared between these four related subjects (Identical by Type). Five SNPs were found to be shared only by both tumor samples. On the other hand, twenty one polymorphisms were shared by P26 and P39 control samples.

### SNPs Identity and sharing

No marker was identified as IBD or HBD between samples. Chi-square association and permutation testing for marker sharing failed to show significant P-values at any marker. The Mantel haplotype sharing statistic using adjusted step-down p-values did not show any significant haplotype sharing around markers (although these findings relate to the small number of tested samples).

### Homozygosity

tumor samples P17 and P61 showed homozygosity at 100% and 97% of markers respectively. Sample P39 showed 29% homozygous SNPs and sample P26 had 23% homozygous SNPs. The observed difference was found significant between tumor samples and related control samples (Monte Carlo simulations p-value < 0.0001 at 0.05 significance level). Significant difference between homozygous SNPs percentages between each tumor sample and control sample was shown using Marascuilo procedure. Runs of homozygosity were seen in Colorectal cancer samples. In sample P17, twenty six consecutive homozygous SNPs were seen in the MHC II region (6p21.32). Samples P61 had twenty five consecutive SNPs in the MHC II region. In both related control samples, P26 and P39, the number of consecutive homozygous SNPs did not exceed 7. Figure (1) and supplementary (1) show these runs of homozygosity in tumor samples.

### Validation by comparison with The Cancer Genome Atlas

We used TCGA data portal to query somatic mutations in colonic adenocarcinoma tumor samples and identified a patient harboring a similar HLA-DQB1 genotype (TCGA ID: TCGA-G4-6298). The patient had four HLA-DQB1 missense SNPs (rs1130390, rs1071637, rs1063318, and rs1049069). The patient was of *Black or African American* race diagnosed with carcinoma caecum stage IIIb at 90 years of age. He had no family history of colonic adenocarcinoma and no synchronous/metachronous malignancies. There was no available data on microsatellite instability, kras/braf mutation status or immunohistochemistry for MMR proteins.

### Predicting the effect of observed mutations

To predict the effect of non-damaging non-synonymous SNPs on the protein, we visualized the position of those variants on canonical HLA protein 3D models. Positions of four variants present only in both tumor samples were visualized on HLA-DQB1 protein structure (figure (3)). All were located in close proximity to the binding groove. On the other hand, All SNPs seen in controls (whether exclusively or shared with tumors) were found to be located relatively distant from the binding groove of HLA-DQB1. Two coding SNPs were predicted to be deleterious on protein structure using ConDel. An HLA-A polymorphism resulting in G80R amino acid change (rs1059449) was predicted to be deleterious on two transcripts (2 and 201) of the HLA-A gene (gene has 10 transcripts). iMutant predicted decreased stability (DDG score −0.52; SNP largely destabilize the protein if DDG<-0.5 Kcal/mol). It was found as a homozygous polymorphism in tumor sample P61. Another polymorphism in HLA-DQA1 gene (rs10093) was predicted to be deleterious resulting in a Q30E amino acid change in gene transcript 008 (gene has 10 transcripts). iMutant predicted weak effect on protein stability (DDG scores of −0.09; SNP has a weak effect if −0.5<DDG<0.5). The two tumor samples (P17 and P61) had homozygous polymorphism, while control samples P26 and P39 had heterozygous polymorphism. Search for variations on TFBS using SNPNexus predicted that rs1063318 affect TFBS for the transcription factor p300. This variant (rs1063318) was mapped to both tumor samples. TFSEARCH confirmed the presence of a binding site for p300 that is affected by rs1063318 but with a slight change in score. On the contrary, Alibaba did not predict a p300 binding site. RegulomeDB scores showed only three variants (rs12363, rs1049053, and rs1064594) with relatively high score, all mapped to controls. No variants were predicted to affect miRNA binding.

**Figure (3).**
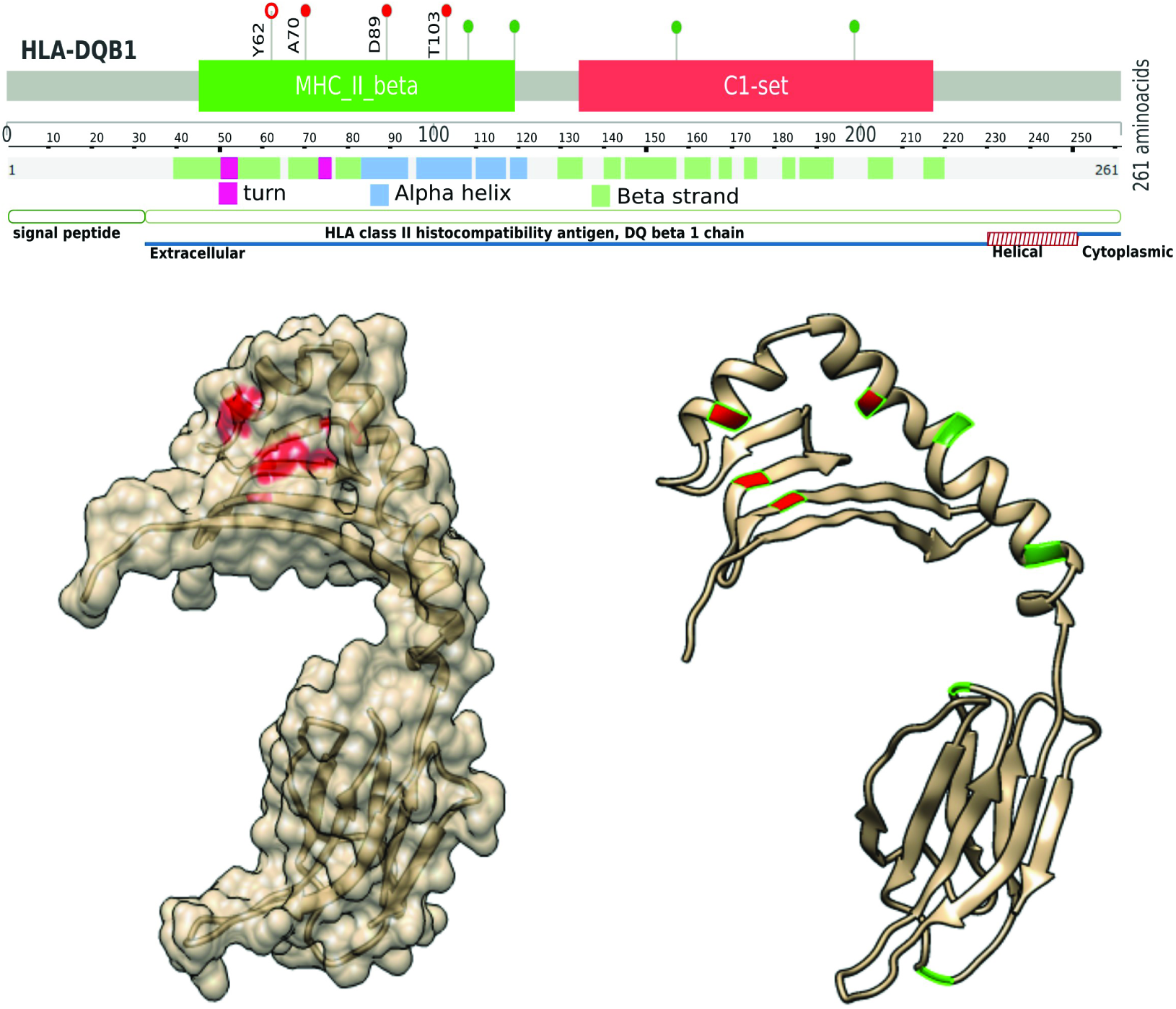
Identical by state SNPs affecting HLA-DQB1 binding groove. The three dimensional structure of HLA-DQB1 protein (ribbon and hydrophobicity models on the right and left respectively) is shown along with the secondary structure of the protein. The position of aminoacids affected by *identical by state* non-synonymous coding SNPs is highlighted in red (in the lollipop plot and 3D model). These SNPs are shared by tumor samples only and affect the HLA-DQB1 binding groove. Positions of other aminoacids affected by exonic nonsynonymous SNPs in tumor and control samples (green) are shown for comparison. MHC_II_beta: major histocompatibility complex class II beta 1 region. C1_set: Immunoglobulin-like C1-type domain.

## DISCUSSION

The role of MHC in colorectal cancer is not well understood. HLA variants have been linked to many types of malignancies including cervical, lung, breast and other malignancies [5-9]. However, a search for phenotypic association of SNPs seen in this study with colorectal cancer did not return a previously reported association. In fact, only a small number of studies, if any, have shown a relation between HLA alleles and CRC risk. A previous study reported that HLA-DQA1* 0201 was less common in 80 Italian patients with colonic carcinoma than controls (8% vs. 18%, P-value with Bonferroni correction = 0.027; OR = 0.44) [6]. Nonetheless, antigen presentation pathway was found to be central in colorectal cancer pathogenesis and showed enrichment in relapsed tumors [30]. Systems Biology analysis also featured a central role for HLA proteins [30]. In this study, identical by state sharing of HLA SNPs between tumor samples in HLA-DQB1 was noted (figure (2)).

High degree of homozygosity was seen in tumor samples in MHC II area (figure (1)). This Run of Homozygosity seen in the MHC II is likely to represent Loss of Heterozygosity (LOH). Runs of Homozygosity (ROH) are not unusual in cancer genomes and in colorectal cancer specifically. Ozaslan and Aytekin have shown LOH in colorectal cancer genomes at multiple chromosomes [31]. Generally, ROH might indicate copy-neutral loss of heterozygosity or copy number variation and subsequent haploinsufficiency. In fact, haploinsufficiency was found to drive aneuploidy patterns and shape the cancer genome. The vast majority of sporadic tumor suppressors are likely to be haploinsufficient [32]. High-frequency LOH was also found to correlate with high metastatic potential of colorectal cancers [33]. These events seem to be somatic in origin. Siraj et al. studied the genome homozygosity in Saudi consanguineous families with CRC family history and concluded that there is no correlation between autozygosity in germline genome and CRC risk [28]. Interestingly, studies did not report LOH at MHC region on chromosome 6p. Although LOH is a common phenomenon in a variety of human cancers, a high frequency allelic loss at a specific chromosomal region indicates the location of a candidate tumor suppressor gene and micro-deletions. In this study it’s premature to conclude the presence of a tumor suppressor locus based on current data alone without considering the frequency of heterozygosity loss. Yet, such a high degree of homozygosity points to this possibility. The increased homozygosity was not seen in control samples.

The central feature of HLA proteins in biological processes remains their ability to recognize and bind peptides. The binding specificity of HLA molecules is basically determined by variations in the surface of the binding groove. The fact that all the non-synonymous coding variants seen exclusively in tumor samples here are in close proximity to the binding groove of HLA-DQB1 emphasizes an active role for the observed haplotype in colorectal cancer pathogenesis (in fact three of them were localized on the surface of the binding groove; figure (3)). This spatial position of shared variants was unique to tumor samples: all the shared variants in controls were distant from the binding groove.

In this study, the observation that SNPs seen in MHC II area are not predicted to be damaging (to most of transcripts) possibly relates to an active role for MHC in CRC pathogenesis and negative selection of damaging variants, making completely detrimental variants an unlikely occurrence. This role is unlikely to be mediated by a damaging effect on protein structure as we argued. For surface MHC I, cells failing to express MHC I induce immune response, at least in early tumor transformation, targeted to eliminate such cells for example by Natural Killer cells [34]. In this sense, immune-editing in colorectal cancer is thought to occur at level of expression of classic and non-classic HLA genes as shown by some studies [35]. However, stable class II MHC is constitutively expressed only in Antigen Presenting Cells surface. Almost all normal gastrointestinal epithelial cells lack surface MHC class II. Gastrointestinal cancer cells often do not express MHC class II molecules, which are associated with T-cell-mediated anti-tumor immune responses [36,37]. Several reports suggest that forced expression of MHC in cancer cells leads to loss of tumorigenicity [38]. Non-viral tumors frequently lose expression of HLA molecules such as the reduction or total loss in colorectal carcinoma [39]. This raises another question on how do these observed variants in MHC II participates in cancer pathogenesis. They are more likely to represent a tumor-selected haplotype that participates in pathogenesis and immune response to cancer. How this can be achieved is likely beyond a classical explanation of surface expression of MHC, especially when we consider the lack of inducible surface expression of MHC II in malignancies. One possibility is that the role of MHC II expression in CRC relates to exosomes and extracellualr vesicle transport. Exosomes are rich in MHC II proteins and have been demonstrated to carry certain markers in cancer [40]. They are also demonstrated to induce tolerance by communication with the dendritic and T-cells in many models. It is plausible as well that exosomes bearing antigen/MHC complexes transmit death signals that cause specific killing of the T cell clones that pose a threat to the exosome-producing cell [41,42].

## CONCLUSION

In this family study, an in-silico analysis was carried out on the results of whole exome capture of two colorectal cancer tumor samples and two control blood samples from healthy related individuals. MHC II area showed high degree of homozygosity in tumor samples evidenced by the presence of runs of homozygosity. This might represent a possibility of disease predisposing or a tumor suppressor locus affected by neoplastic transformation. Shared markers were found non-identical by descent. No marker or haplotype sharing was proved statistically significant using non-parametric tests of significance, but this might be explained by the rather small number of studied samples. Sharing of three exonic non-synonymous variants was seen between tumor samples. Tumor-shared SNPs, unlike those of controls, were predicted to affect the binding groove of HLA-DQB1 protein and thus affect its binding specificity. The results demonstrate IBS SNP sharing and possible loss of heterozygosity in classic MHC II region in cancer tissues. The significance of this sharing is yet to be determined. Changes in binding specificity and extracellualr transport seem to be plausible explanations for HLA role in this study.

### Supplementary Data

Supplementary (1) is a figure depicting unphased haplotypes of single nucleotide variations seen in MHC area in this study, detailing the positions, sharing, and homozygosity.

### List of Abbreviations

CRC: colorectal cancer. GERP: Genomic Evolutionary Rate Profiling. HBD: Homozygous By Decent. HLA: human leukocyte antigen. HNPCC: Hereditary Non-Polyposis Colorectal Cancer. IBD: Identical By Decent. IBS: Identical By State. LOH: loss of heterozygosity. MHC: major histocompatibility complex. MMR: Mismatch Repair. ROH: run of homozygosity. SFVTs: sequence feature variant type sites. TFBS: transcription factor binding site. TCGA: The Cancer Genome Atlas.

### Authors' Contributions

MK, SS, MI: contributed to the conception and design. SS, MS, MI: participated in data collection. MK, MA: carried out the analysis. MK, MA, MI: wrote the primary draft. All authors revised the manuscript critically for intellectual content and approved the final version.

### Authors' Information

Mahmoud E. Koko (Institute of Endemic Diseases, University of Khartoum), Suleiman H. Suleiman (Soba University Hospital, University of Khartoum), Mohammed O.E. Abdallah (Institute of Endemic Diseases, University of Khartoum), Muhallab Saad (Institute of Endemic Diseases, University of Khartoum), Muntaser E. Ibrahim (Institute of Endemic Diseases, University of Khartoum, 11111 Khartoum, Sudan).

### Ethics statement

This study is an in-silico analysis of open-access published data that is cited inside manuscript. The study was ethically approved by the ethical and scientific committees of the Institute of Endemic Diseases, University of Khartoum. Written informed consent was obtained from all participants, in accordance with the Declaration of Helsinki.

